# Antibiotic-mediated interactions underlying microbial diversity

**DOI:** 10.1101/2023.02.15.528676

**Authors:** Gaurav S. Athreya, Chaitanya S. Gokhale, Prateek Verma

## Abstract

The immense diversity observed in natural microbial communities is surprising in light of the numerous weapons microbes have evolved to inhibit each other’s growth. It is thus imperative to understand which interaction patterns can sustain a biodiverse community when individual species antagonistically affect one another. In this study, we leverage potent methods from theoretical ecology to show how antibiotic-mediated interactions between microbes drive biological diversity. Building on previous experimental and theoretical results, we analyse the dynamics induced by various interaction graphs involving antibiotic production, resistance, and degradation. Previous work has recognised the importance of a particular producer-sensitive-degrader (PSD) motif in the interaction graph. We study this motif in detail and elucidate the mechanistic reason for this importance. Concretely, we give exact rules for coexistence in some simple cases where exhaustive enumeration of the interaction graphs is feasible. More generally, our results suggest that the PSD motif, in combination with a cyclic interaction structure, is sufficient for stable coexistence in well-mixed populations. Using individual-based simulations, we then study the importance of the PSD motif in spatially structured populations. We show that community coexistence is robust for an extensive range of antibiotic and degrader diffusivities. Together, these findings illuminate the interaction patterns that give rise to diversity in complex microbial communities, stressing that antagonism does not imply a lack of diversity and suggesting clear approaches for culturing synthetic microbial consortia.

## Introduction

Studies of various microbial ecosystems have shown that microbial communities can harbour immense biodiversity [1, 2, 3]. In particular, there has been empirical evidence from studies of pairwise interaction networks that competitive interactions might be more prevalent than cooperative interactions [4]. In light of the competitive exclusion principle [5], the existence of high microbial biodiversity is non-intuitive in such communities. Therefore, we focus in this work on how microbial communities are stable in the face - or perhaps because of - interference competition. While there are many types of weapons that microbes have evolved, we consider chemical weapons due to their prevalence and the rich history of research on antibiotic-mediated interactions [6, 7, 8]. An antibiotic, hereafter, is any compound secreted by a microbe that leads to an increase in the death rate of other microbes either by killing them (bactericidal) or by stalling cellular processes that prevent further growth (bacteriostatic). These antibiotics inhibit other microbes by affecting varied processes like core metabolism, the cell envelope, or transcription-translation machinery. Conversely, it has also been shown that these “weapons” can have varied functions such as signalling and growth stimulation when present in low concentrations and are, therefore, not always responsible for inhibitory interactions [9, 10, 11].

The strains in microbial communities produce and are resistant or sensitive to several antibiotics [12, 13]. Moreover, the pathways that enable these properties are also modular, with genes often found on plasmids, making them prone to horizontal gene transfer [14, 15]. Since each strain can have a different phenotype concerning each antibiotic, numerous community configurations are possible. Empirical studies have investigated the interaction networks of soil-dwelling microbes and found many properties of these networks that are non-trivial and distinct from other ecological networks [16, 17].

In this context, we are interested in finding interaction graphs that give rise to communities that support species diversity even in the in the presence of antagonistic/competitive interactions. Early work on the effect of interspecific interactions on population dynamics showed that cyclic antagonistic interactions between three species could lead to a cyclical dominance in a well-mixed setting [18]. However, this coexistence is unrealistic since abundance is assumed to go arbitrarily close to zero, which is impossible since individuals are discrete. This discrete nature will result in stochastic extinction events after a sufficiently long time [19]. However, in a spatially structured population with an identical interaction graph, coexistence has been observed via the formation of patches that chase each other around the lattice and via moving spirals that are lost as dispersal radius increases [20, 21]. Therefore, spatial structure can play an essential role in determining biodiversity.

Interactions mediated by antibiotics are a specific type of antagonistic interaction. Subsequent studies have shown that coexistence is possible in spatially structured communities with antibiotic-mediated interactions [22, 23, 24]. However, understanding how such communities can remain diverse, especially in unstructured environments, has been a significant challenge due to the strong inhibitory nature of antibiotic chemicals. Kelsic et al. (2015) [25] showed robust co-existence in a community with cyclic antagonistic interactions where all antibiotics are attenuated by a degrader phenotype. Degraders secrete chemicals that inactivate the antibiotic, so they confer resistance to themselves and other individuals in their neighbourhood. The degradation of antibiotics is known to be a common mechanism of resistance [26, 27].

In particular, Kelsic et al., (2015) [25] consider a cyclical interaction graph with three strains with each strain being a producer, degrader and sensitive for one of the three antibiotics (see Figure 1). This interaction structure was prevalent in the communities that they found in soil-dwelling bacterial communities. They show that this configuration is stable in both the well-mixed and spatially structured population cases. Degradation allows fine-scale intermixing of the three strains, as opposed to large-scale patches, as previously observed with cyclical interaction graphs [28, 20].

**Figure 1:**
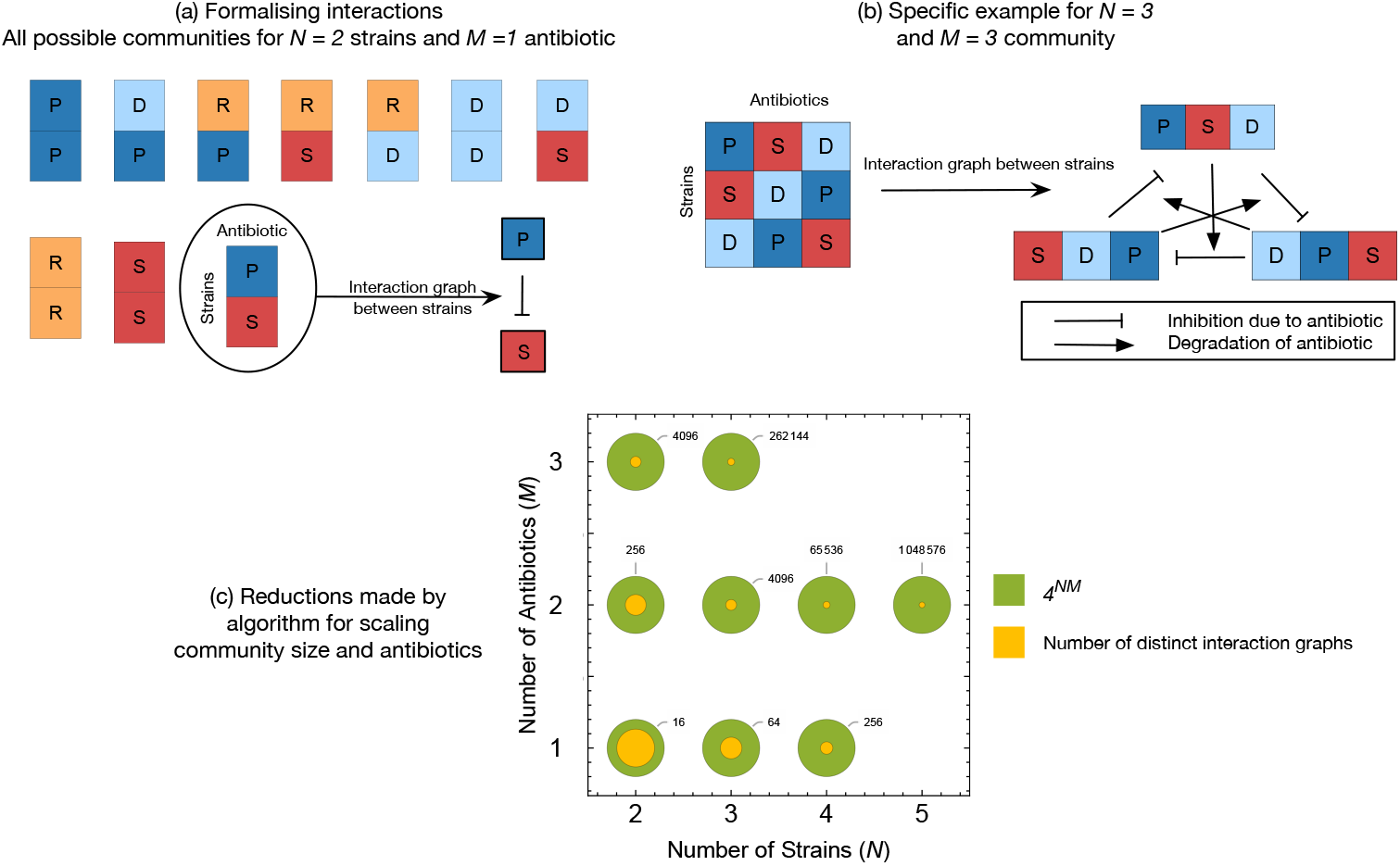
Enumeration and analysis of model microbial communities. **(a)** We formalise the interspecies interaction graphs by assuming that the phenotypes are discrete - an individual can either be a producer(P), sensitive(S), degrader(D), or resistant(R) with respect to a given antibiotic. This panel shows the ten possible distinct communities with *N* = 2 strains and *M* = 1 antibiotic. **(b)** The interaction graph of the community analysed in depth in [25]. Our study examines more generally the conditions necessary and sufficient for the existence of stable states in such communities. Any interaction graph corresponds to a set of equations, described in section 2.1. We provide a method to generalise this analysis to communities of any *N* and *M*. Our strategy is to search for stable steady states in all possible communities with a fixed number of strains (*N*) and antibiotics (*M*), and then to vary *N* and *M*. Thus checking for all possible interaction graphs, this procedure allows us to infer the exact conditions for stability. **(c)** In section A.1 of the supplementary information (SI), we develop and describe an algorithm to enumerate all interaction graphs with a fixed number of strains and antibiotics. The naive expectation for the number of unique interaction graphs with *N* strains and *M* antibiotics is 4^*NM*^ (outer circles) since there are four possible phenotypes - P, S, D, R, but this is untrue (see section A.1, SI). This figure compares this naive expectation with the actual number of distinct interaction graphs (inner circles) for different values of *N* and *M*.

Given our knowledge of the diversity and complexity of interactions in real communities, it is natural to ask whether the above coexistence mechanism translates to more extensive and complex community assemblies. Kelsic et al. (2015) [25] have also shown that increasing the number of antibiotics that can be produced or degraded by the community leads to richer dynamics with the ability to support more diversity. However, the larger communities explored in their work were assembled by inexhaustive random sampling all possible strain combinations that can make up a community, and their results hence provide only broad-level statistics. Hence, a systematic study of the particular conditions required for the ecological stability of more complex communities, identifying the motifs of interactions that lead to stability, is necessary but lacking. In this work, we first provide results for stability in communities with a range of complexities, which allows us to characterise the conditions required for stability in three-strain, three-antibiotic communities. Secondly, we study the effect of spatial structure on the stability of a community with one antibiotic and three strains - a producer strain, a sensitive strain, and a degrader strain. This simple community exhibits stability [29]. Here, we examine the effect of chemical diffusivities on this community’s population dynamics and show that its stability is remarkably robust to changes in these parameters.

## Model and Results

### 2.1 Ecological dynamics: the mixed-inhibition zone model

Much theory in ecology assumes pairwise interactions, but the chemicals secreted by microbes can indirectly affect other strains [30, 31]. The widely used classic Lotka-Volterra formalism cannot adequately describe these higher-order interactions [32]. Therefore we generalise and use the mixed-inhibition zone model introduced by Kelsic et al. (2015) [25]. We assume well-mixed populations and discrete phenotypes - a given strain can be a producer (P), degrader (D), intrinsically resistant (R), or sensitive (S) to a particular antibiotic.

Production, intrinsic resistance, and degradation are metabolically expensive, with the highest cost for production followed by degradation and then intrinsic resistance. This cost structure is assumed because producers both produce antibiotics and are resistant to them, degraders, secrete chemicals and imbue several individuals with resistance, whereas intrinsic resistance benefits only one individual. Assuming a constant base fitness for all strains, producers have the least growth rate and sensitive strains the largest. Let *c*_*p*_, *c*_*d*_ and *c*_*r*_ be the costs of antibiotic production, degradation, and intrinsic resistance. Note that we make the strong simplifying assumption that the costs for each antibiotic are the same. This is obviously unrealistic, but its benefits are twofold - besides simplifying calculations, it allows us to isolate the effect of interaction graphs on population dynamics. When we consider multiple antibiotics, the base growth rate of a particular strain is reduced in an additive fashion depending on how many antibiotics a strain can produce, degrade, and so forth.

Our model has two types of chemicals - the antibiotics themselves and the chemicals used by the degraders to degrade the respective antibiotic. These chemicals can diffuse around the individuals secreting them. To formalise this, we assume, following [25], that there is a critical region around an individual in which the chemical is effective and ineffective outside. In the next section, we shall consider a more realistic case, where we explicitly model diffusion by incorporating a gradient around each producing/degrading individual.

Consider a microbial community composed of *N* distinct strains. Each strain can be a producer *P*, degrader *D*, resistant *R* and sensitive *S* with respective to *M* different antibiotics. Let *P, S* and *D*, and *R* be *N* × *M* matrices that store the phenotype of each strain concerning each antibiotic. The entry *P*_*ij*_ = 1 if strain *i* is a producer of antibiotic *j*, and the other matrices are defined similarly. Note that for a given index (*i, j*), exactly one of *P*_*ij*_, *S*_*ij*_, *D*_*ij*_, *R*_*ij*_ is 1 and all others are zero. If *g* is the base growth rate of strain *i*, then the effective growth rates *r*_*i*_ are given by,

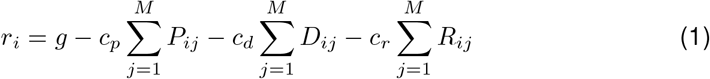

Next, we find the probability 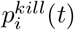 of an individual of strain *i* dying due to the action of antibiotics at time *t*. Note that this value must be nonzero precisely when strain *i* is sensitive to antibiotic *j* (*S*_*i,j*_ = 1) and zero otherwise. We can find the probability 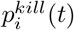 of dying (due to at least one antibiotic) by using the inclusion-exclusion principle. Let 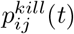 be the probability of a strain *i* individual dying due to the action of antibiotic *j*. Then we have

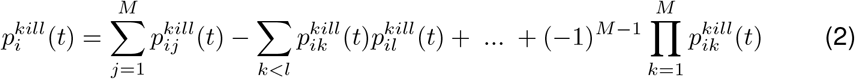

where the direct summation of the 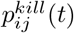 is incorrect since these events are not mutually exclusive - an individual of the focal species can simultaneously be inhibited by two or more antibiotics that it is sensitive to.

Now we calculate 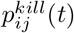 for a given strain *i* and antibiotic *j*. To be killed by antibiotic *j*, an individual of strain *i* must be sensitive to it, be outside all degrader zones and inside the production zone of at least one producer. Therefore we must calculate the probability of being outside and inside these zones respectively. Let *K*_*P*_ and *K*_*D*_ be the effective killing area around each producer and the effective degradation area around each degrader. We use the Poisson process on the plane as a model for individuals and their (random) position on the Petri plate. If *λ* is the density parameter for this process, the probability of finding zero points in a region of unit area is given by *e*^*−λ*^.

The abundance *X*_*i*_ can be interpreted as the fraction of the area occupied by strain *i*. But the production (resp. degradation) zones of antibiotic *j* may arise due to individuals from any strains producing (resp. degrading) this antibiotic - of which there are at most *N*. Therefore, we sum over the *j*th column of the matrices *P* and *D* to find the total abundance of producers and degraders of antibiotic *j*. Consequently, the fraction of area covered by production zones for antibiotic *j* is 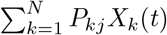 (and similarly 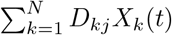 for total degradation zone area). The density *λ* is replaced as applicable by the above quantities, i.e., the area fractions in which production and degradation are effective, respectively. Of course, if strain *i* is not sensitive to antibiotic *j*, then 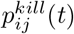 must be zero for all *t*. Hence, we have

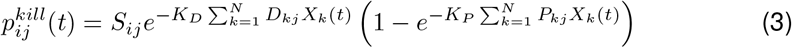

Using equations 2 and 3, we can then write the fitness *f*_*i*_ of a strain at time step *t* as

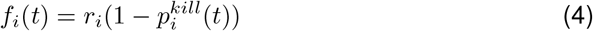

as based on the growth and death rates of the strains [33]. We use a discrete version of the replicator equation [34] for the population dynamics

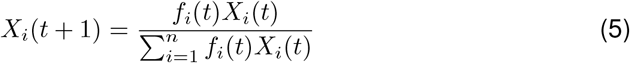

As an illustration of the model, consider the community {P, S, D} with three strains (*N* = 3) interacting via one antibiotic (*M* = 1) (Figure 2(b), left). Let us index the strains in the same order. The growth rates of these strains and probabilities of dying due to antibiotic action are given by,

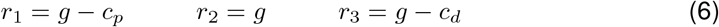

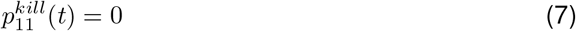

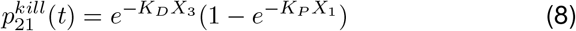

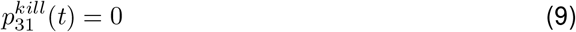

which can be plugged into Eqs. (4) and (5) describing the complete community dynamics.

**Figure 2:**
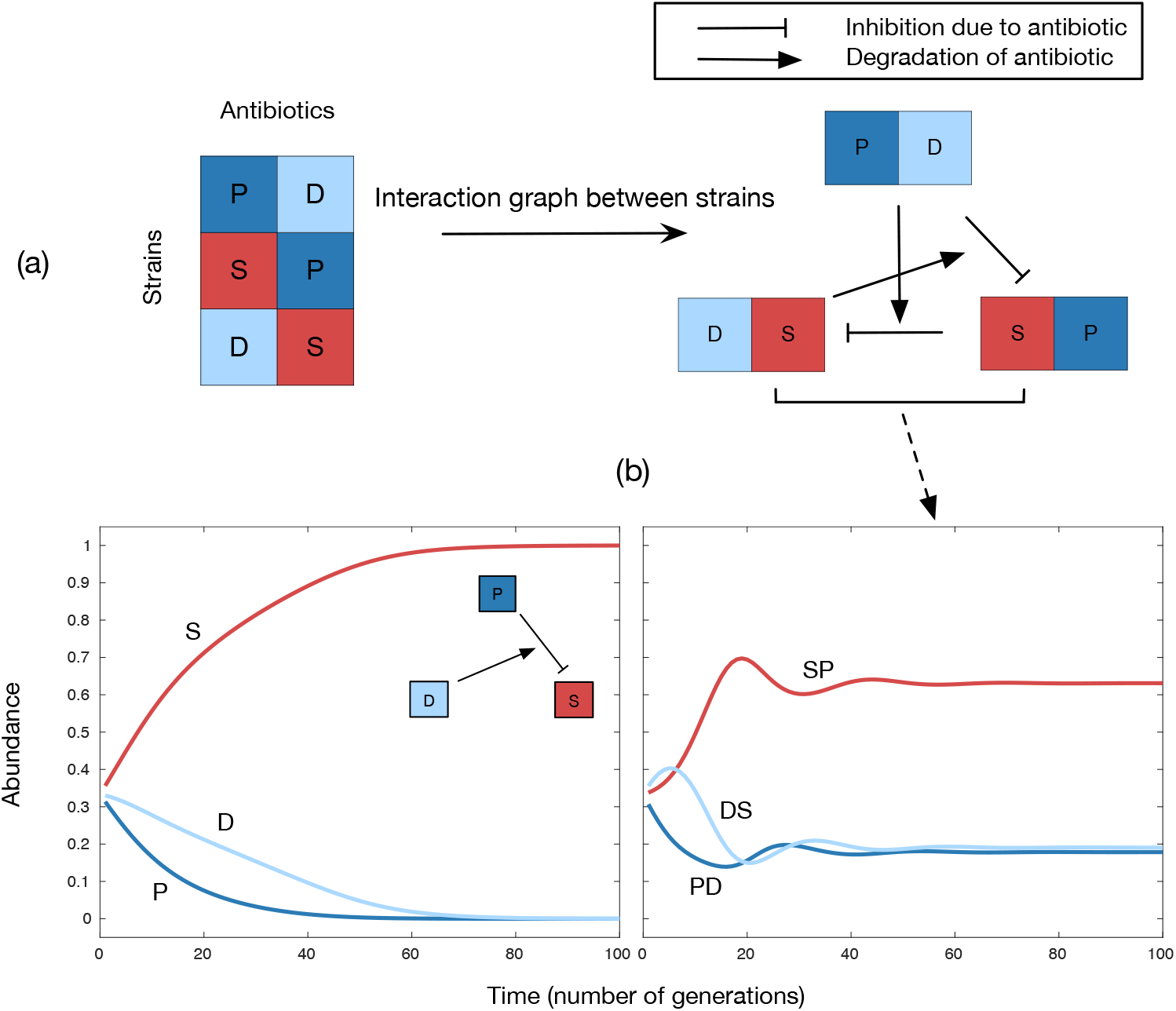
Existence of a minimal complexity for stable co-existence. We check for the existence of a stable fixed point in several pairs of *N* and *M* values. Of these, *N* = 3, *M* = 2 is particularly interesting since it is the first combination for which there is a stable fixed point, i.e., it contains the minimal complexity required for a stable internal fixed point to exist [25]. But this complexity is insufficient - the interaction structure must also be appropriate. Using our exploration framework Sec. 2.2.1, we show that the only stable interaction structure with *N* = 3, *M* = 2 for the parameter combinations we have checked is the one displayed in part **(a)**. It is, therefore, important to understand which properties of this pattern are essential in this respect. **(b)** We have a “PSD motif” concerning a particular antibiotic when a Producer, Sensitive, and Degrader strain of that antibiotic is present in that community. Consider a *N* = 3, *M* = 1 community with a PSD motif and the *N* = 3, *M* = 2 community pictured above. The dynamics of the three strains in these two communities is qualitatively different - in the left panel, the sensitive strain (S) wins, whereas in the right panel there is stable coexistence. We argue that this is because the different antibiotics can have - as in this particular community - opposite effects on the population dynam12ics. The runaway S strain in the left plot is therefore “controlled” due to the dynamics induced by the second antibiotic. Parameter values: *g* = 1.0, *c*_*p*_ = 0.15, *c*_*d*_ = 0.105, *c*_*r*_ = 0.05, *K*_*P*_ = 10, *K*_*D*_ = 15, initial abundance = 1/3

### 2.2 Results and analysis of the mixed-inhibition zone model

We shall now use this model to analyse communities with different numbers of strains and antibiotics. The strategy will be to explore, for several *N* and *M* values, the space of all communities with *N* strains and *M* antibiotics and determine whether (and if so, which) communities with that many strains and antibiotics can exhibit stable states.

The space of all unique communities having *N* strains and *M* antibiotics can be enumerated efficiently by the algorithm outlined in section A.1 of the Supplementary Information (SI), the results of which are available in Fig. 1. Our objective is to search for communities with stable internal fixed points, which correspond to a steady-state supported with non-zero abundances of all involved strains. We find these fixed points and determine their stability by numerically solving a system of equations for each community having those *N* and *M* values. In particular, a fixed point is stable if the maximum absolute value of the eigenvalues is less than 1.

It is, of course, possible for populations to coexist without stable fixed points via, for example, limit cycle oscillations; more generally, see the ideas of persistence and permanence in population dynamics [35, 36]. Here we focus on stable fixed points for their analytical and computational tractability.

#### Notation

We will denote communities by a vertical array of *N* rows (i.e., a matrix), with each row denoting a strain in the community. The rows have *M* letters from P, S, D and R, and the *j*th letter signifies the phenotype of the given strain with respect to the *j*th antibiotic. In the text, such a community will be denoted by a list of its rows. For example, the community on the right in Figure 1 is denoted by *{*PDS, DSP, SPD*}*.

#### 2.2.1 Exhaustive enumeration of interaction graphs

The equations of the mixed-inhibition zone model simplify in different ways depending on the interaction structure in a particular community, as seen in the given example. We numerically solve these equations for each community arising from the (*N, M*) values as given in Figure 1. Explicit enumeration allows us to give the exact conditions required for stability since we check all possible interaction graphs. To our best knowledge, there is no closed-form expression for how the number of distinct interaction graphs scales with *N* and *M*, so enumeration becomes a non-trivial task for higher complexity communities. See Table 1 for a full record of these results.

**Table 1:** For multiple combinations of *N* and *M*, we looked at the ecological dynamics induced by all possible interaction graphs for those many strains and antibiotics. A record of *N* -*M* combinations for which there exists a stable internal fixed point is shown in this table. Note: The number *N* here refers to the *distinct* number of strains.

| N (number of strains) | M(number of antibiotics) | Stable internal fixed point present |
| --- | --- | --- |
| 2 | 1 | No |
| 3 | 1 | No |
| 4 | 1 | No |
| 2 | 2 | No |
| 3 | 2 | Yes (unique) |
| 4 | 2 | No |
| 5 | 2 | No |
| 2 | 3 | No |
| 3 | 3 | Yes (several) |

**Table 2:**
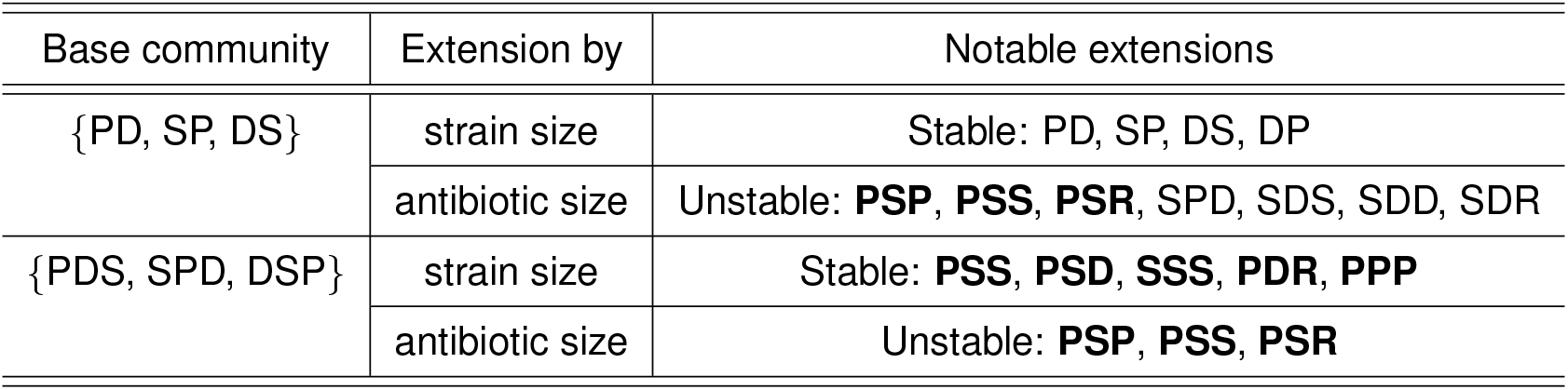
We focus on two base communities which we know have robust stable fixed points at the parameter combination we are working with. We extend the interaction graph of these communities in two ways: either by adding another strain (another row in previously developed notation) which can interact via existing antibiotics, or by adding another antibiotic (another column) through which existing strains can interact. For each combination of base community and type of extension, we give an exhaustive list of notable extensions that lead to either loss or maintenance - as specified in the table - of the stable fixed point. All parameter values are kept unchanged when adding new rows or columns. For example, stable extensions are those where the addition of that row or column does not destroy the stable fixed point and hence preserves stability. Unstable extensions are defined analogously. **Emboldened**: all permutations of these extensions are stable/unstable as well. Parameter values: *g* = 1.0, *c*_*p*_ = 0.15, *c*_*d*_ = 0.105, *c*_*r*_ = 0.05, *K*_*P*_ = 10, *K*_*D*_ = 15.

We begin by looking at the case of *M* = 1, i.e., one antibiotic. Under the assumptions of this model, we observe that for *N* = 2, 3, and 4, there are no stable internal fixed points. ^1^ In particular, the communities {P,S,D} and {P,S,R} do not have any stable fixed points - recall that the results of [29] and [23] show that spatial structure induces stability in these communities.

Note that in all cases below, a community composed of *N* identical phenotypes is “degenerate” in that any initial condition is a fixed point - there can be no nontrivial dynamics if all the *f*_*i*_ are the same (see Eq (5)). Similarly, a community of *N* strains, 2 of which have identical phenotypes, has a similar degeneracy - the effective number of strains is *N −* 1, resulting in a family of solutions. These cases arise merely as an artefact of our looking at all possible communities. In the results that follow, these degenerate cases are implicitly excluded when considering stable internal fixed points.

For *M* = 2, no community with two strains has a stable internal fixed point, i.e., all communities reach a single-strain composition after a sufficiently long time. A producer strains can lose to a sensitive strain if the metabolic cost of production and the initial abundance of the sensitive strain is sufficiently high.

When *N* = 3, however, we find that precisely one of 430 distinct communities has an isolated stable internal fixed point, and this is the *{*PD,SP,DS*}* community, shown in Figure 2. ^2^ This community has a producer, degrader, and sensitive strain (henceforth called a “PSD motif”) corresponding to each antibiotic. Recall that the results from [25] and [29] suggest that this condition might be important for stability. Importantly, for *N* = 3 and *M* = 2, there is more than one way to have a PSD motif with respect to each antibiotic, but the only interaction graph that induces a stable state is the one where interactions are made to be as cyclic as possible (see Figure 2, as opposed to, for example, *{*PS, SP, DD*}*. This combination of *N* and *M* is also the minimal case for which there can be a stable internal fixed point.

Notice further that there is no stable community with 4 or 5 strains and 2 antibiotics. This suggests that at least in well-mixed populations, the conditions for stability are very strict and specific interaction graphs are necessary. Issues of computational power limit us from searching for the next largest community on 2 strains that has a stable community.

When the community has 3 antibiotics, the minimal number of strains required for stability is 3. For *N* = *M* = 3, there are several communities with a stable internal fixed point. One of these is of course the community analysed in detail by Kelsic et al. [25]. The other communities are easy to characterise: they all contain PSD motifs with respect to at least two antibiotics.

But it has not yet been explained why there is a minimal complexity required for stability - with the PSD motif on one antibiotic, the sensitive strain wins for a large number of *K*_*P*_ and *K*_*D*_ values. This is due to the assumption of well-mixed populations and the interplay between antibiotic action and metabolic costs. However, with two antibiotics, the impact can be opposite in effect - a strain can increase in abundance due to the action of one antibiotic and decrease due to another. This is precisely what happens in our system (see Figure 2) - the sensitive strain that always used to win now also has an additional phenotype due to which its abundance is driven down. The interplay of these dynamics leads to the existence of a stable fixed point. However, this balance is, in a sense, not exact with two antibiotics since the phenotypes are not all symmetric. They are symmetric, however, in the case of the community analysed by Kelsic et al. with 3 antibiotics. We see an exact balance there, evidenced by the equal abundances of 1/3 of all strains (the abundances are not 1/3 in the case of 2 antibiotics). For symmetry (or lack thereof) of phenotypes in the PSD communities, see the interaction graphs of these communities in Figures 2 and 3 (left). In summary, the results of this exhaustive enumeration, some of which have been obtained already in [25], suggest that the PSD motif is vital for community stability.

**Figure 3:**
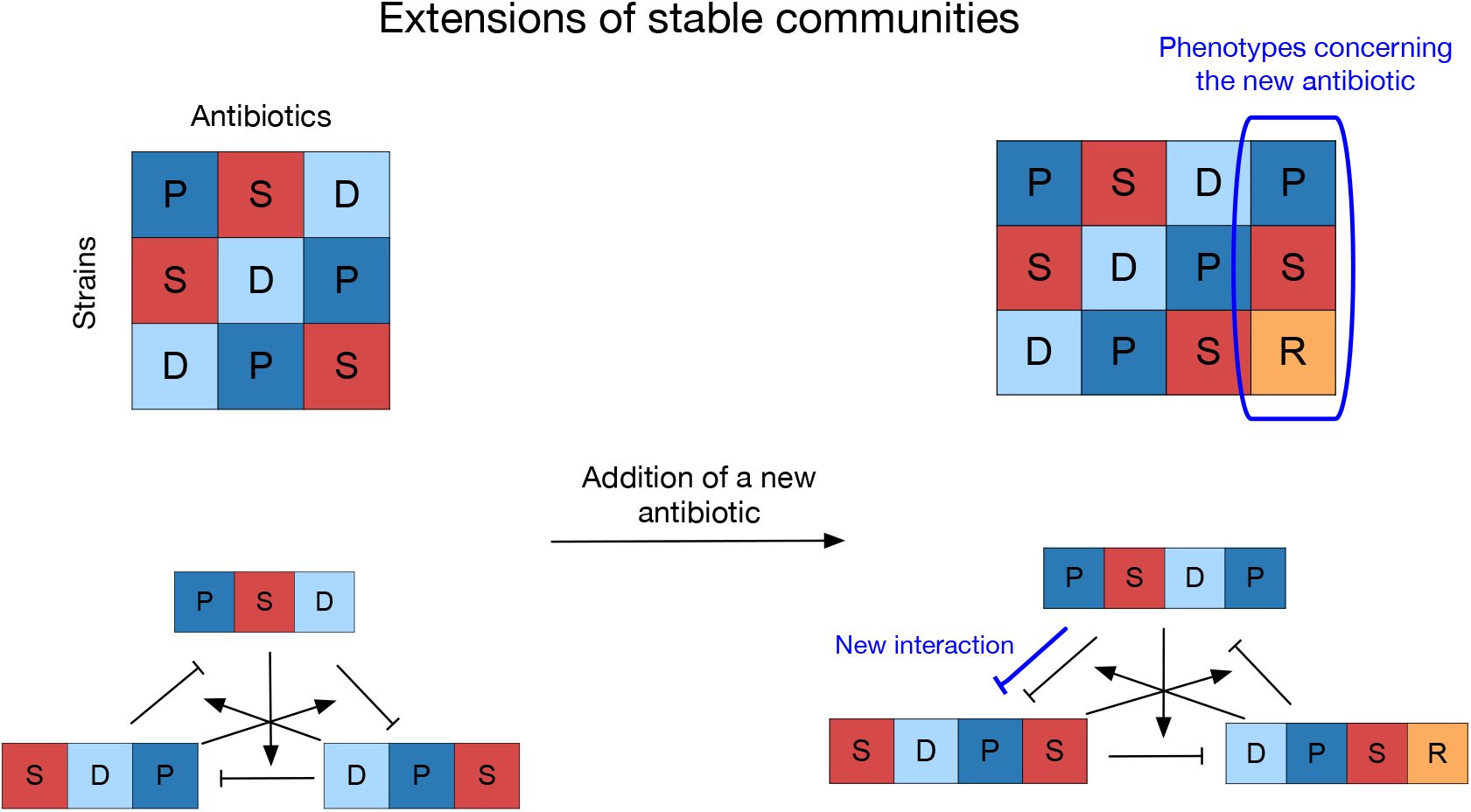
A community in which the strains of base community 2 (left) have evolved the ability (highlighted, right) to produce and resist a new antibiotic. These kinds of “extended” communities are helpful to consider for two reasons. Firstly, the extensions coarse-grain the microevolutionary processes in these communities and help us understand their properties analytically. Secondly, they represent a simple way to study how more complex patterns might allow for more diversity. We show that in some cases, only a particular kind of extension preserves the existence and stability of the fixed point - there has to be a stabilising interaction (or lack of a destabilising one) concerning each antibiotic. Particularly, degradation is the only way to stabilise a producer-sensitive inhibitory interaction in this model.

#### 2.2.2 Extensions of stable communities

In Nature, microbial communities are in constant flux - both ecologically and evolutionarily. Consider the scenario when a community, through evolutionary processes, has reached an ecologically stable state. However, note that the model in [16] predicts that microbial communities in some cases are never in an ecologically stable state and are constantly in a state of flux with rapid turnover of antibiotics and strains, but this view does not seem to be widely held. Such communities can be perturbed via further ecological destabilisation of ongoing evolutionary dynamics [37, 38]. In particular, this could take the form of immigration of new individuals into a stable community or mutation of resident phenotypes, which can correspond to an increase in the number of strains *N*. Alternatively, resident strains can evolve production and resistance capabilities to a new antibiotic - corresponding to an increase in the number of antibiotics *M*. Another source of motivation for such an analysis is trying to understand the existence of exactly which communities are stable in the case of *N* = *M* = 3. For *M* = 2 there is exactly one, but the increase of antibiotics by 1 induced stability in many communities - which are these communities, and how are they related to this unique *N* = 3, *M* = 2 stable community?

To understand how such increases in complexity affect the stability of a community, we consider two simple communities that contain a stable steady-state. The communities from Figure 2 (base community 1), and Figure 1 (base community 2) [25], both of which already exhibit the PSD motif. In line with the motivation detailed above, we increase these communities’ *N* and *M* (for computational tractability, only by 1). We then exhaustively search the set of all communities with interaction graphs that can be obtained by such an extension.

First, we distinguish between two types of phenotypes concerning the new antibiotic. Consider the representation of this extension in Figure 3 - in the additional column, a producer-sensitive pair either exists or does not. If it does not, all strains resist the new antibiotic since S is the only sensitive phenotype, and sensitivity is binary in this model. Then, the additional column only amounts to a decrease in the growth rates of the strains and does not introduce a qualitative change in the dynamics. However, if a producer-sensitive pair exists, it has a definite destabilising effect since the strain that produces the new antibiotic inhibits the strain that is sensitive to the new antibiotic.

First, let us consider the addition of antibiotics. We find that if a new column contains a producer-sensitive pair for *{*PDS, SPD, DSP*}*, the stable fixed point is maintained if and only if the third strain degrades the new antibiotic. The exact coordinates will, of course, be different due to changes in growth rates since they produce, degrade, or resist an additional antibiotic. There must exist a “PSD motif” concerning the new antibiotic. This observation supports that degradation is the only mechanism stabilising producer-sensitive pairs in our well-mixed model. Furthermore, all communities where the new column does not contain a producer-sensitive interaction still have a stable fixed point. - a fact that we think asserts the robustness of these stable fixed points to variation in growth rates.

In comparison, a statement similar to that for the earlier base community holds with an essential caveat for the base community 1 *{*PD, SP, DS*}*. Here as well, all producer-sensitive interactions without a degrader are unstable. The caveat is that there are some more communities where the stable fixed point is lost. Further, enumerating *N* = 3, *M* = 3 communities with stable fixed points shows that the only *N* = 3, *M* = 3 communities to have a stable fixed point are those where all producersensitive interactions are degraded (see associated GitHub repository). This shows that the PSD motif is sufficient for stability but not necessary.

The “extra” unstable extensions can be partitioned into two types in terms of what we interpret them to mean. First, the triplet SDS, SDD, SDR - notice that this is the set of all extensions which start with SD excluding SDP. SDP is somehow the only extension which preserves stability. Note that SDP is precisely the extension required to give base community 2, which has both the PSD motif and whose strains are symmetric (giving rise to an utterly symmetric interaction graph). This shows that if the phenotype is partly imposed (by specifying “SD_”), the only way to preserve stability is to have the extension that makes the strains symmetric. It may be asked why this does not occur when other parts of the symmetry-inducing extension (“SDP”) are incompletely specified, such as S_P and _DP. The first is easy - that S_P is only compatible with D has already been established since it has a producer-sensitive pair. However, it is less clear for _DP; any of the 4 phenotypes in the empty place lead to a stable extension. Whatever the correct interpretation, the general point emerges that symmetry and cyclicity of interactions are also important in addition to the PSD motif. We are left now with the unstable extension SPD. Note that all other SPD permutations are stable-PSD, PDS, SDP, DPS, DSP. This again shows that the PSD motif is not sufficient but is crucial. In general, we conclude that the importance of the PSD motif is still evident. Still, the asymmetry in the strain phenotypes of base community 1 leads to the extra complications required for a stable extension. We note again that the exact conditions and stable extensions might change with changes in parameter values. Still, the general message will likely stay since the base communities’ fixed point is robust.

Now we consider the addition of strains. This is motivated by the biological scenario where strains of another phenotype immigrate into a stable community, or new phenotypes evolve. Instead of starting at small migrant abundance, we focus on the question of whether or not there exists a stable fixed point in this “augmented” community. As long as the abundances are inside the presumed fixed point’s basin of attraction, the new strain can stably coexist with the resident strains. For base community 1, the first 3 - PD, SP, DS - are just duplicates of strains already present and are hence trivial, and there is another stable extension - DP. For base community 2, **PSD** contains all duplicates and some more stable extensions. PSS, SSS, PDR show that strains with very different kinds of phenotypes can coexist with the base community. Unfortunately, there does not seem to be an easy characterisation of the stable extensions here. However, the classes of non-trivial stable extensions are again more in number for base community 2, and we interpret this to be because of (the lack of) symmetry.

In conclusion, these findings suggest that the degradation of antibiotics, and the “PSD motif” in particular, might be a fundamental force of stabilization in microbial communities with antibiotic-mediated interactions. It suggests that the independent means of interaction (each antibiotic) must together have a balanced structure. Further, we find that the base community with more antibiotics and symmetry is harder to destabilise in multiple ways.

### 2.3 Ecological dynamics in spatially structured populations

Kelsic et al. [25] used a 3-species, 3-antibiotic, 3 degrader enzyme system to show that multi-species coexistence is possible even in well-mixed populations regardless of attempts at invasion by cheater mutants. However, the impact of the degrader on antibiotics was taken into account by considering the formation of circular inhibition zones of fixed radii within which all antibiotic is degraded and all sensitive species protected. A more realistic treatment should account for diffusion of both degrader enzyme and antibiotics in a systematic manner with degraders being able to neutralise the effect of antibiotic closer to their location and less capable of doing so farther from their location. In view of this drawback, it is useful to analyse a realistic model of interaction between degrader, sensitive and antibiotic producer species. Further, previous literature provides strong evidence that the complicated conditions required for stability (as above) may be restricted to well-mixed models and that the class of stable communities in a spatially structured setting is much larger [20, 23, 29]. Therefore, it is instructive to study spatially structured populations to understand the effect of local interactions and diffusive processes on community stability. As a first attempt to understand this presumably larger class of stable communities, we study in detail one of its simplest members - the simplest possible PSD motif. This motif has 3 strains and 1 antibiotic, and we find that it does not exhibit a steady state over a large part of the parameter space in the mixed-inhibition zone model. Moreover, as an instance of the PSD motif, this community’s behaviour might lead to insights into how the rules that hold in the well-mixed case transform under the addition of spatial structure. It has, however, been shown to be stable by Vetsigian [29] in a slightly different implementation. In this section, we summarise a spatial model of communities with antibiotic-mediated interactions with a detailed analysis of motif stability as a function of the parameter values. For a detailed description, see Supplementary Information, Section A2. Note that the construction of this model does not use any specific properties of the PSD motif and can thus be applied to an arbitrary community.

The lattice is initialised first by deciding the occupation of a given site with probability *p*. If a site is chosen to be occupied, then the species identity of the individual that occupies it is chosen uniformly randomly from 1, …, *N*. At each time step, we iterate over all sites on the lattice. If the focal site is empty, we move on. If it is occupied, we induce secretion of ay relevant antibiotic or degrader chemicals as applicable depending on the phenotype of this individual. Then the chemicals diffuse, and we keep track of the cumulative concentrations at each site. Then, we consider the birth and death of this individual. An individual dies with a probability that depends on the antibiotic concentration on the site that it occupies, and the cell is then empty. An individual gives birth only if there is an empty neighbouring site (of which there are 6), and the precise location of the offspring is chosen uniformly from these neighbours. This procedure is then iterated for a long time, until we are confident that an attracting dynamical object (fixed point, cycle, etc.) has been reached.

### 2.4 Results and analysis of the stochastic spatial model

As is observable from the previous section, individual-based models allow for realistic considerations without the need for complicated mathematical machinery. While our model can be used to simulate any community with *N* strains and *M* antibiotics, we use it to study the simplest case where the minimum complexity for coexistence is revised directly due to the presence of spatial structure, i.e., in the 1-antibiotic PSD motif. Recall that previous simulations have shown that the *{*P,S,D*}* motif can be both ecologically and evolutionarily stable [29].

#### 2.4.1 Spatial structure strengthens the PSD motif

Consider the 3-strain, 1-antibiotic community consisting of a producer strain, a sensitive strain, and a degrader strain. We study the ecological stability of this community as a function of the diffusivities of the antibiotic and degrader chemicals. To do this, we simulated the population dynamics of the spatial model over an extensive range of these diffusivities, given by *K*_*P*_ and *K*_*D*_ values respectively. *K*_*P*_ and *K*_*D*_ set two aspects of the model’s behaviour - one is the width of the Gaussian filter; the other is the shape of the dose-response curve. The parameters *K*_*P*_ and *K*_*D*_ do not change the metabolic cost since they are the diffusivities - which only depend on the environment and the amount of chemical produced by each individual is kept constant. In particular, 15 independent simulation runs were carried out, each 6,000 generations long, for each of 900 parameter combinations, ranging from *K*_*P*_ and *K*_*D*_ values between and 18. This is chosen to keep *σ*_*P*_ > 1, where *σ*_*P*_ is the standard deviation of the Gaussian that controls the spread of the antibiotic chemical. This was imposed to find some constants in the functional form of the dose-response curve. More specifically, a high value of *K*_*P*_ corresponds to a large diffusivity and a higher susceptibility to the antibiotic (due to the expression of the above constants in terms of *K*_*P*_, see SI for details of the spatial model). Some examples of parameter values for which the population dynamics relaxed onto a stable state are shown in Figure 4.

**Figure 4:**
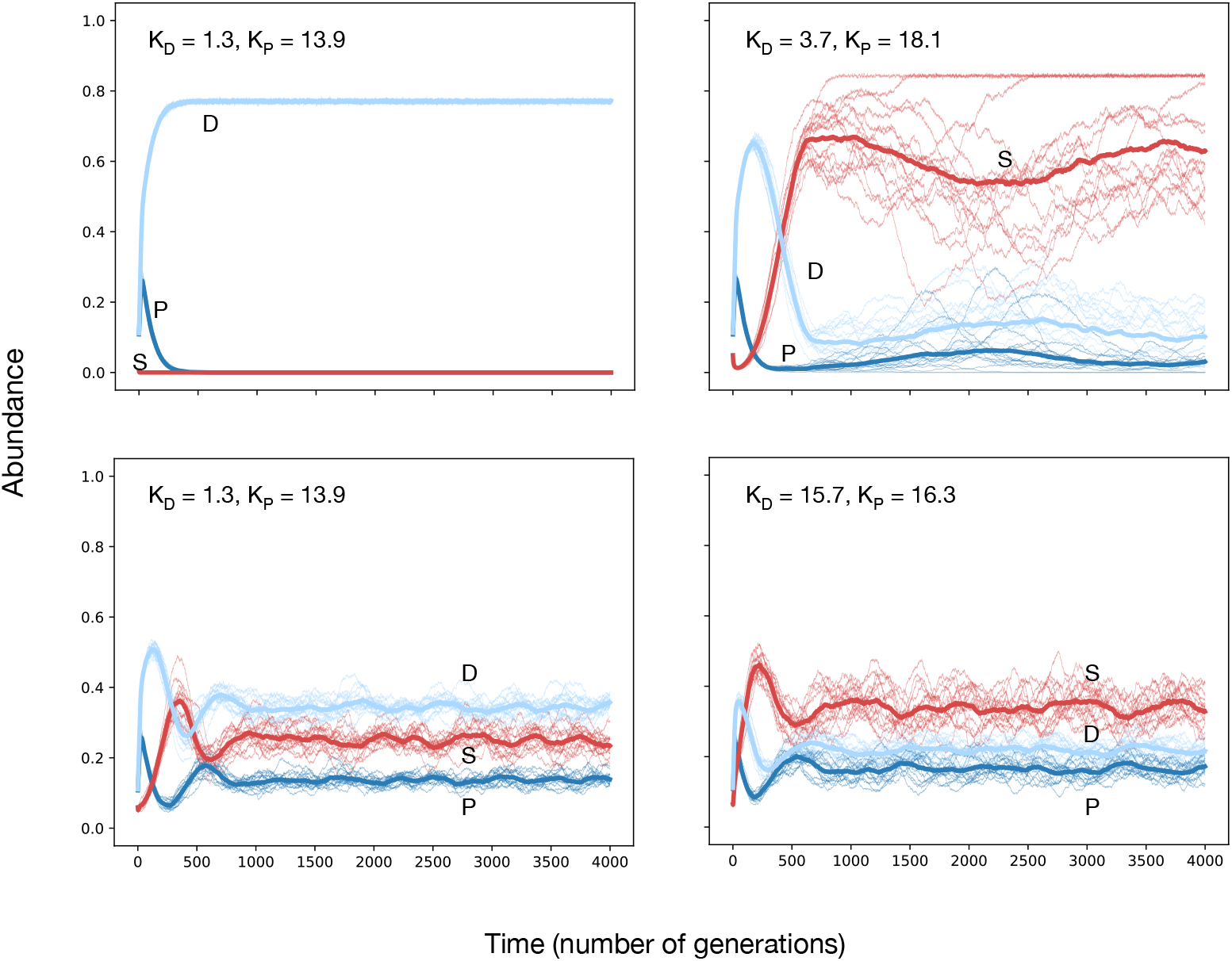
Spatial structure induces coexistence in the 1-antibiotic PSD motif. We run individual-based lattice simulations with periodic boundary conditions on a community with three strains - a producer, sensitive, and degrader. The lattice is initialised with an occupancy probability of 0.3 per grid cell, with the identity of an occupied cell chosen at random. For each parameter combination, 15 independent simulation runs were carried out, each 6,000 generations long, for each of 900 parameter combinations, ranging from *K*_*P*_ and *K*_*D*_ values between 0.7 to 18. Therefore, the initial configuration of the lattice is different in each run, and only the occupation fraction is kept constant. Outside of intrinsic birth and death, the individuals interact via chemicals that they produce or are sensitive to. These chemicals diffuse over the lattice and this induces long-range higher-order interactions. Here we show representative examples of some parameter combinations for which there is robust coexistence. From the trajectories, we see that there is a stable steady state for these parameter combinations, with small fluctuations around this steady state. Note here that the relative abundances is a function of the diffusivities. Figure 5 shows the composition of the population at the steady state over a much larger range of *K*_*P*_ and *K*_*D*_ values. Parameter values: *g* = 0.7, *d* = 0.3, *c*_*r*_ 2=1 0.05, *c*_*d*_ = 0.105, *c*_*p*_ = 0.15, grid size = 200, *u*_*p*_ = 10, *u*_*d*_ = 10

Figure 5 shows the result of these simulations over the explored *K*_*P*_ *−K*_*D*_ parameter space. The colour of the grid cell for a particular pair of *K*_*P*_ and *K*_*D*_ values denotes the average population abundance over the last 500 time steps of the simulation. The abundances can be identified using the simplex on the right as a legend.

**Figure 5:**
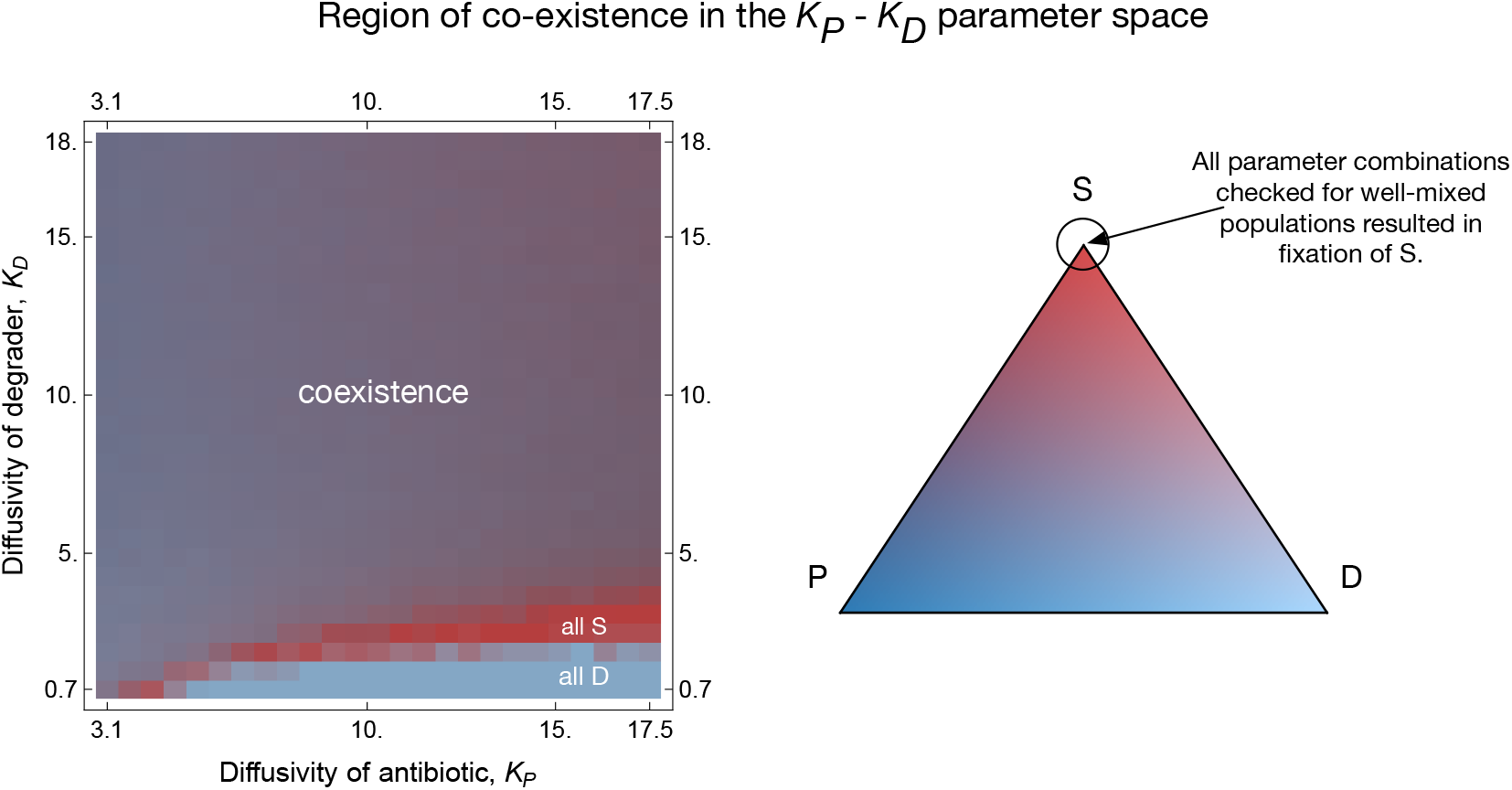
Ecological dynamics in the *K*_*P*_ *− K*_*D*_ parameter space. We analyse the ecological dynamics over a range of diffusivities of the antibiotic and degrader chemicals, presented as a heatmap on the left. The simplex on the right serves as a legend for the colours in the heatmap in the following sense. Each point on the simplex corresponds to a set of three numbers *P, S, D* such that *P* + *S* + *D* = 1, which we interpret as population abundances. For a given value of *K*_*P*_ and *K*_*D*_, the colour of the heatmap cell, through this interpretation of the simplex, therefore shows how many producer, sensitive, and degrader individuals were present on the lattice at the end of the simulation. The vertices of the simplex are “pure” colours and correspond to all S (red), all P (dark blue), and all D (light blue) lattices. Other colours are combinations of the three pure colours and correspond to a population with coexisting types. The dynamics of well-mixed PSD populations is in sharp contrast to spatially structured populations - for all *K*_*P*_ and *K*_*D*_ values checked, the sensitive strain always grew to fixation. However, the population at steady state is composed of near-equal abundances of all three strains over an extensive range of the parameter space. The *K*_*P*_ and *K*_*D*_ of the lattice and well-mixed model are not identical. The unidimensional strength of the chemicals in the mixed-inhibition zone model is split into two notions of strength in the spatial model. The amount of the antibiotic produced and its diffusivity jointly determine the extent to which surrounding individuals experience the antibiotic. They are, however, positively correlated since diffusivity determines overall strength. Parameter values same as Figure 4.

In Figure 5, we compare the state of the population after a long time under different combinations of *K*_*P*_ and *K*_*D*_. The first thing one might notice is the large region of coexistence between all three strains - this implies that the attractor is robust to changes in diffusivities. Further, there is a clear boundary along which coexistence is lost to fixation by the sensitive strain, below which the degrader strain fixes. Although perhaps trivial, it is essential to note that the mere existence of spatial structure cannot stabilise populations at all parameter values. This happens for the following reason - for large *K*_*P*_ and small *K*_*D*_, the sensitive strain is killed off very quickly due to high diffusivity of the antibiotic and the producer is then outcompeted by the degrader strain because of its lower metabolic cost. For slightly higher *K*_*D*_, the degrader strain can protect the sensitive strain enough to prevent its extinction. Simultaneously, the degrader outcompetes the producer, after which the sensitive strain outcompetes the degrader. In other words, the deciding characteristic is the *timescale of local interaction* between patches of different phenotypes. Thus, we have shown that this community’s ecological stability is very generic, especially when contrasted with the lack of stable states in the mixed-inhibition zone model.

#### 2.4.2 Existence and nature of the boundary

Next, in more detail, we analyse Figure 5 to understand its nature. The structure suggests a separatrix - the coexistence equilibrium above it (green), fixation of the degrader below it (blue), and the fixation of the sensitive strain along the boundary between these two regions (red). The dynamics on the boundary can be observed in movies of the simulations (see associated online repository). Along this boundary, we observe that the sensitive strain wins in a significant fraction of the cases (see Figure 4, top right). In some other simulation runs, the degrader strain wins.

We claim that these dynamics can be entirely understood in the following way. The producer strain is always outcompeted to begin with, and the eventual fate depends on whether or not a patch of sensitive individuals has persisted long enough for the producers to have died out. If these persistent, sensitive individuals still exist, they outcompete the remaining degraders based purely on the difference in growth rates. If the sensitive strains have been killed off by the producers, the degraders can win without impeding the sensitive individuals’ superior growth rate. Therefore, along this boundary, *K*_*P*_ is weak enough to not kill the sensitive individuals in the initial generations and the growth rates of such that the producers are killed off in these same initial generations. The persistence (or lack thereof in some cases) of the sensitive strain then determines the long-term fate of the population. For *K*_*P*_ any higher, the producer can kill off all the sensitive strains, and the degrader then wins. For *K*_*D*_ any higher, the sensitive strains can be protected against antibiotics, and the usual cyclical dynamics take over. Therefore, the boundary is characterised precisely by the fixation of the sensitive strain. We expect this boundary to collapse into a one-dimensional curve in a deterministic model. Any departures from that expectation in our results are due to the finite-size stochastic nature of our individual-based model. Based on the above explanation, we expect the location of this boundary to move with changes in the metabolic cost but for the shape to stay similar. This suggests a single steady state exists in this system’s interior, which could be recovered in a deterministic model using systems of partial differential equations for the abundance of strains and concentration of the antibiotics. Given the current study’s scale, this promising yet extensive research avenue is deferred to the future.

## Discussion

Antibiotic-mediated interactions are widespread in nature and have been implicated in diverse ecological roles both when produced internally by a microbial community and when present due to external anthropogenic influences [39]. Many microbes produce antibiotics, and medical antibiotics are also often derived from naturally occurring compounds that have been in production long before human intervention [10, 8]. Natural microbial communities have therefore had to stay stable for a long time in the face of many compound ecological effects. When the strains produce these chemicals, and their effects are antagonistic, the abundance of negatively affected strains is driven downwards, and it is therefore unclear how such communities can stay stable. We consider the role of strains that degrade antibiotic chemicals produced by other strains. We also study the patterns in which these strains must produce antibiotics and antibiotic-degraders for a stable, co-existing steady state for the community population dynamics.

The interactions between the strains in a community can be formalised in terms of mathematical objects called graphs. We describe a strategy, limited only by computational power, to exhaustively analyse the dynamics induced by a wide range of interaction graphs. These graphs can be precisely enumerated using the algorithm described in section A.1 of the Supplementary Information. Following Doebeli et al., (2017) [33] and Kelsic et al. (2015) [25], we derive fitnesses rooted in birth and death rates derived mechanistically from the strain phenotypes. The ecological dynamics specified by each pattern are then analysed numerically for the existence of a stable fixed point. This procedure allows one to make a complete record of the behaviour of all interaction graphs up to a complexity feasible given the available computational power. With this in mind, we ask a simple question: which interaction graphs give rise to a stable fixed point for the community’s population dynamics?

Previous studies of similar systems have identified a particular motif in the interaction graph, the producer-sensitive-degrader (PSD) motif, that is potentially important for community stability [25, 29, 40]. Concretely, a PSD motif is said to occur concerning a particular antibiotic in the interaction graph if at least one strain produces the antibiotic, a strain that is sensitive, and a strain that degrades it. Recall that a strain can have only one of these phenotypes with respect to an antibiotic. Firstly, Kelsic et al. analyse the properties of a 3-strain, 3-antibiotic community in the strains that have phenotypes such that there is a producer, sensitive, and degrader strain concerning every antibiotic. Theoretical work on spatially structured populations [29] has shown that allowing the production and degradation of one antibiotic can give rise to a 3-strain community (producer, sensitive, and degrader strains) that is both ecologically and evolutionarily stable. Doing the same with two antibiotics shows that there is the formation of a stable 5-strain community - and that the composition of this stable community has a producer, degrader, and a sensitive strain for both antibiotics [29]. The potential importance of this motif is clear from these observations, and we, therefore, study a simplified model from Kelsic et al. [25] to obtain a mechanistic understanding of how and when this motif is essential.

We find that, for a small number of antibiotics, there are simple rules that can ensure the existence of a stable state for the community dynamics. The PSD motif is indeed essential, but it is not so trivial as to say that it is merely necessary and/or sufficient for co-existence. There are a few caveats to such statements that our work identifies, and we shall state these first before proceeding to the main conclusion. Firstly, cycles in interactions or the presence of a PSD motif themselves are not sufficient for stability in well-mixed populations. This is evidenced by the observation that the counterpart PSR motif does not exhibit a stable fixed point despite having a cyclic interaction graph. The PSD motif is not sufficient since *{*P, S, D*}* has a PSD motif but no steady state. What is clear is that degradation of all producer-sensitive interactions seems to be sufficient when there is more than one antibiotic - we have also explained why multiple antibiotics are required for stability (see Figure 5). This claim is equivalent to the condition of an existence of a PSD motif for every antibiotic for which there is a producer-sensitive pair. However, these PSD motifs must balance each others effects on the strains - this is inherent to the explanation in figure 2 and gives rise to the cyclic structure in the interaction graphs. Therefore, it is a combination of the PSD motif and cyclicity in the interactions that leads to stable co-existence. This is the fundamental, mechanistic reason why the PSD motif has a stabilising effect. For example, {PP, SS, DD} technically has two PSD motifs - one with respect to each antibiotic - but does not have a stable fixed point. In contrast, {PD, SP, DS} has two PSD motifs in a cyclic structure, which leads to a stable fixed point. In conclusion, PSD motifs are important, but not unconditionally, and the mechanism of stability inherently draws on a variety of antibiotics through which the strains might be interacting. This is especially encouraging since the strains in natural communities are likely to produce and degrade many chemicals.

We must, however, temper this with the knowledge that cyclic dominance has been shown to be rare in evolving Lotka-Volterra communities [41]. Our work suggests that even though these communities might rarely arise, cyclic dominance, when present with PSD motifs, is a strong force for stabilisation even in communities with higherorder interactions. In this sense, we observe modularity in the interactions of these communities that emerge in the long run. Our work further confirms that this property of interaction graphs is indeed important by exhaustive enumeration and can lead to co-existence even if it is not present at the level of each antibiotic.

Further, we have shown that the interaction graph influences the stability of a coexisting community to the evolution of new antibiotics or the arrival of new strains. The base community with more antibiotics and symmetry was harder to destabilise, and it would be interesting to see in future work how the complexity of a community affects invadability or susceptibility to the evolution of additional antibiotics.

However, note that allowing different diffusivities for different antibiotics, which is more realistic than the simplification we have made, might change these rules. For example, a low-diffusivity antibiotic causing instability because of a producersensitive interaction which is not degraded may not lead to the removal of a fixed point if there is another high-diffusivity antibiotic for which the producer-sensitive interaction is degraded.

The method in this study is relevant in the context of migration and founder effects. If a stable community migrates only partly, what fraction of strains is required to maintain co-existence? The analysis of extended communities suggests that a degrader is necessary to stabilise producer-sensitive interactions in our system, and the stable fixed points from our base communities are robust to changes in growth rate. However, note that it is unclear how robust these claims are to the evolution of more antibiotics - this imposes a high metabolic cost, possibly destabilising the steady-state. Alternatively, if some interaction mechanisms in a stable community are removed, how often is the modified community still stable? This last question is especially relevant given the ubiquity of numerous interaction mechanisms (e.g. secondary metabolites can act as growth inhibitors, growth promoters, public goods [6, 16, 42]) in natural microbial communities. Furthermore, the action of these compounds is presumably environment-dependent. Our framework can address such questions by considering extensions and analogous restrictions of the interaction graphs of stable communities and studying fractions of stable communities.

To further understand the validity of this rule in realistic conditions, we also investigate in greater detail the most straightforward instance of the PSD motif in spatially structured populations. Recall that this pattern gives rise to an unstable community in the well-mixed setting. We use individual-based simulations to explore spatially structured population dynamics. Including structure, robust co-existence is possible for a range of diffusivities of the antibiotic and degrader. There exists a smooth boundary between the regions of co-existence and fixation. The degrader strain wins at high *K*_*P*_ and low *K*_*D*_ for specific parameter ranges (see Figure 5). In contrast, the three strains co-exist at all other parameter combinations. Interestingly, the boundary seems to be characterised by the fixation of the sensitive strain. The shape of the boundary and this distinct behaviour along the boundary indicate a bifurcation taking place along these parameters in phase space. This result, in contrast with the well-mixed case, shows that spatial structure is crucial in studying interactions and the possibility of co-existence even with higher-order interactions. Interestingly, controlling the extent of diffusion of the degrader chemical amounts to changing the biological phenotype of the degrader strain. If the degrader chemical does not diffuse or diffuses very little from the source site, degraders, instead of acting altruistically in protecting the sensitive species, behave as loners in which the process of neutralising the antibiotic can be thought to occur within the bacterial cell [43]. Our results also indicate that multispecies co-existence can be environment dependent, with co-existence possible in only those environments allowing for comparable rates of diffusion of the antibiotic and degrader enzymes.

One of the most important aspects of microbial communities is the fast evolution of their constituent strains and the consequent strong coupling between ecological and evolutionary dynamics. This eco-evolutionary feedback is made possible by the short life cycles of the microbes and mechanisms such as horizontal gene transfer. It is very instructive to derive exact rules in simplified models like ours. Nevertheless, it is also necessary for the future to consider how the evolution of antibiotic production/resistance/degradation impacts the eventual community diversity in larger communities [29]. This will require large-scale computational models or conceptual models studying eco-evolutionary dynamics in highly diverse communities with higherorder interactions [44, 45]. Earlier studies have allowed individuals in a community to acquire production/resistance capabilities for as many as 14 different antibiotics. However, these communities did not involve degraders [46, 47]. This community per-spective is also critical in the context of the potential non-attainability of evolutionary stable states [48], and the possibility of inaccessible ecologically stable states [25].

Several more complex factors not included in our model are likely to influence the survival of production and resistance mechanisms in the real world. If the antibiotic had a long lifetime in the environment, this would prevent colonisation by sensitive species of a region previously occupied by producer strains. We also assumed that the production and degradation of the chemicals are rapid compared to the cell division time-scale, so the local antibiotic production is proportional to the local concentration of producers at the current time step. A natural extension of our work would be to relax the aforementioned assumptions and consider the proportional cost of producing antibiotic and degradation enzymes with corresponding increasing strength, the effect of limited nutrients, as well as inter-species conflict and cooperation.

Our findings also suggest better practices in engineering stable microbial communities with desired species. This might have many applications, especially those that benefit from a community perspective. Firstly, many bacteria cannot currently be cultured in the laboratory [49]. Understanding more about the community ecology of unculturable bacteria might be central to improving this situation. The PSD motif we have described is a precise recipe for constructing stable communities and might be an explicit answer for unculturable microbes interacting via antibiotics. This has many applications, one of which is simplifying the screening process for compounds with potential antibiotic effects. Instead of searching for antagonistic interactions, it might be sufficient to search for evidence of the PSD motif, which would then imply the existence of a toxin being produced by a strain of this community. Of course, natural communities are orders of magnitude larger than the ones we have analysed [6]. However, a detailed understanding of small communities has strong translational relevance and is a necessary step in understanding their more complex counterparts.

In conclusion, the outcome of this work is two-fold: it provides intuition for the kind of antibiotic-mediated interactions a stable community can be expected to have. Secondly, we leverage fast numerical computation and an exhaustive search algorithm to obtain some exact conditions despite a complex dynamical system that is hard to analyse analytically. Improvements and modifications of this algorithm can potentially be used to significant effect when considering larger classes of interaction structures. Such methods can only become more important in the future as successive stages of biological reality are introduced into models, making them progressively more intricate and applicable.

## Acknowledgments

Funding from the Max Planck Society is gratefully acknowledged (GSA, CSG, PV). GSA also thanks the INSPIRE-SHE programme of the Department of Science and Technology, Govt. of India.

## Data Availability Statement

All relevant scripts and figure generation pipelines are available at the following public repository: https://github.com/tecoevo/antibiotics_biodiversity.git

## Supplementary information to

## A.1 Removing redundancies in community space

This section describes how the set of all communities with *N* strain and *M* antibiotics (henceforth Ω_*N,M*_) can be efficiently explored. The mixed-inhibition zone model requires the specification of *NM* phenotypes, implemented here by the introduction of the matrices *P, S, D*, and *R*. For the purposes of this section, these 4 matrices can be condensed without loss of information to one *N* × *M* matrix *T*, which has entries *T*_*ij*_ such that *T*_*ij*_ can take values in {1, 2, 3, 4}, corresponding to the possible phenotypes of strain *i* with respect to antibiotic *j* - producer (P), sensitive (S), degrader (D), and intrinsically resistant (R) respectively. See Figure A.1 for an example.

**Fig. A.1:**
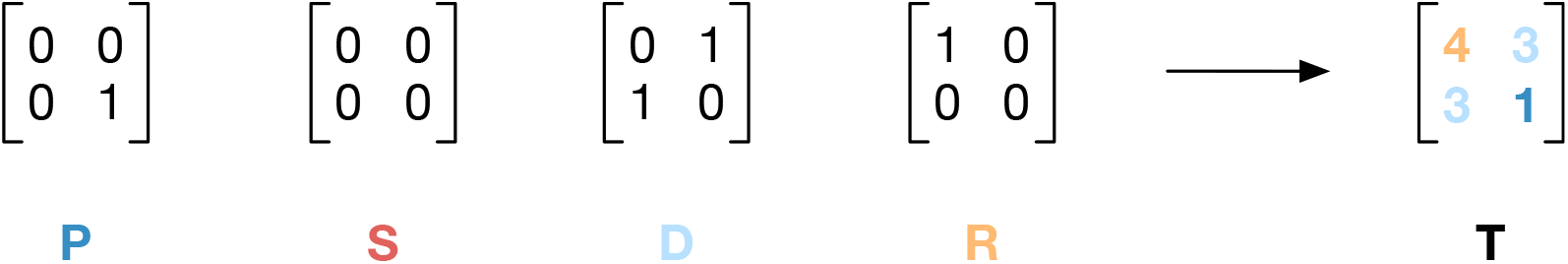
We describe a microbial community in our model and computer scripts by a collection of 4 binary matrices *P, S, D, R* where the letters stand for Producer, Sensitive, Degrader, and Sensitive respectively. These matrices function as follows: *P*_*ij*_ = 1 iff strain *i* is a producer of antibiotic *j, S*_*ij*_ = 1 iff strain *i* is sensitive to antibiotic *j*. However, they can be condensed into one matrix *T*. Here, we show an example for *N* =2, *M* =2 of this condensation of *P, S, D, R* into *T ∈* T_2,2_, the set of all 2 × 2 matrices with entries in {1, 2, 3, 4}

We are therefore describing Ω_*N,M*_ by using the set T_*N,M*_ of all *N* × *M* matrices with entries in {1, 2, 3, 4}. By the assumption of 4 discrete phenotypes, the size of T_*N,M*_ is 4^*NM*^. It is obvious that T_*N,M*_ contains all communities since any community can be described by a set of 4 matrices *P, S, D, R* which can then be condensed to an element of T_*N,M*_. However, the important distinction is that T_*N,M*_ contains multiple copies of each community. The subject of this section is to understand the form of this redundancy and describe a method by which it can be eliminated. It is of course immensely useful to be able to eliminate these copies since naively describing Ω_*N,M*_ using T_*N,M*_ and subsequently performing the analyses on each element of T_*N,M*_ entails a large waste of resources.

As an instructive example, consider the case of *N* = 2 and *M* = 1, that is, the case of 2 strains and 1 antibiotic. T_2,1_ is of size 16, but explicit computation shows that the size of Ω_2,1_ is 10, i.e., smaller than 16. Suppose we describe communities by their strain composition, and we describe each strain using an ordered string of letters from {P,S,D,R} to denote the strain’s phenotype with respect to each antibiotic. The communities in Ω_2,1_ are thus {P,P}, {S,S}, {D,D}, {R,R}, {P,S}, {P,D}, {P,R}, {S,D}, {S,R}, and {D,R}. The 2×1 matrices in T_2,1_, however, take into account the difference in ordering between {P,S} and {S,P} - but a community containing 1 producer strain and 1 sensitive strain is the same as a community containing 1 sensitive strain and 1 producer strain. Doing this reduction by hand becomes increasingly cumbersome with increasing *N* and *M*, and it is therefore useful to have a method of performing this reduction for arbitrary *N* and *M*.

We begin by noting that to interpret the matrices in T_*N,M*_ as encoding strain phenotypes of a community as above, it is necessary to fix a labelling of strain and antibiotics. It is this labelling that induces the redundancy since there is no biological significance to it - when interpreting an element of T_*N,M*_, we infer from it the phenotype of each strain with respect to each antibiotic, but a given strain has the same phenotype irrespective of whether we call it strain 1 or strain 2 (similarly for the antibiotics). Consider for example the two pairs of matrices shown in Figure A.2. The first pair has rows interchanged - denoting a different labelling of the *N* strains, whereas the second pair has columns interchanged - denoting a different labelling of the *M* antibiotics. Notice that all 4 matrices, despite being different elements of T_2,3_, describe the same community.

**Fig. A.2:**
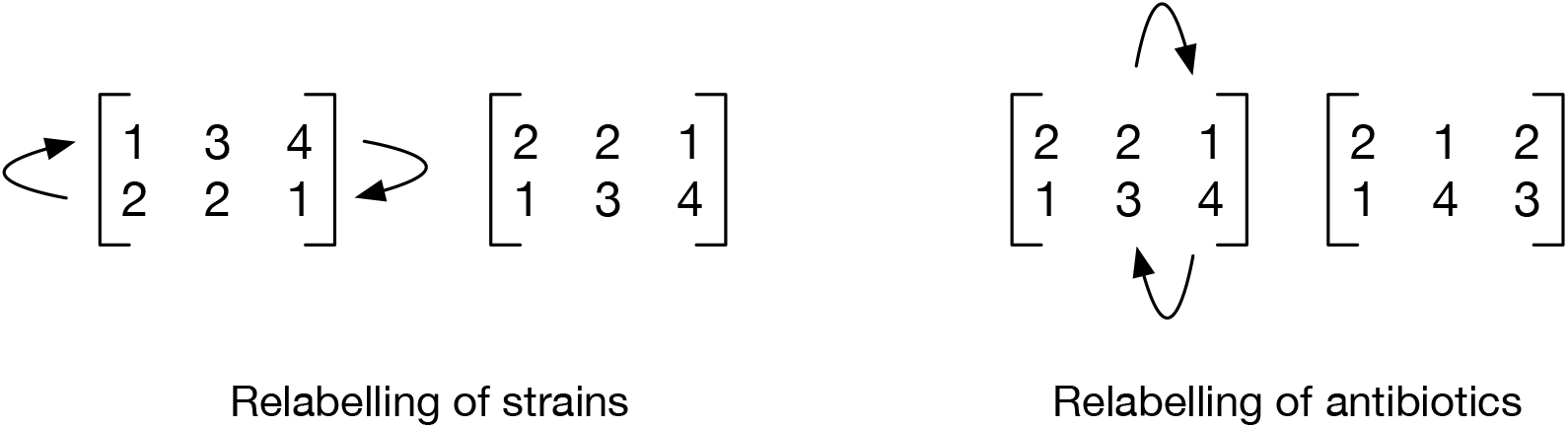
The matrix description even after condensation as in the previous figure is redundant - the identity of a matrix is not uniquely given by a matrix in T_*N,M*_. Instead, many matrices describe the same community. This is because a community does not change if the strains or antibiotics are merely relabelled - called 1 and 2 (in order) instead of 2 and 1. This order is externally imposed, and so does not change the community. This figure shows redundant description of the same community by different elements of T_2,3_.

The redundancy described so far can be otherwise understood as follows: a community is identical under changes in how the strains and antibiotics are labelled. The labelling is given by the order of rows and columns in the matrices of T_ℕ,𝕄_, and so communities are identical under permutations of rows and columns. In other words, despite an arbitrary number of permutations of its rows and columns, a matrix will describe the same community. It is therefore natural to turn to the idea of graph isomorphisms since the structure of a graph is also identical under different labelings of its vertices and edges.

With this intuition, we remedy this redundancy as follows. Let *T* be a matrix in 𝕋_*N,M*_ and let *C ∈* Ω_*N,M*_ be the community described by *T*. First we transform *T* into a labelled bipartite graph which we will call the *antibiotic profile* of *T*, with the two parts representing the strains and antibiotics. This transformation on the elements of 𝕋_*N,M*_ gives rise to a set 𝔾_*M,N*_ of bipartite graphs. We then perform a reduction on 𝔾_*M,N*_ using the notion of an isomorphism of graphs, and convert the reduced set of graphs back into matrices.

Given *T ∈* 𝕋_*N,M*_, construct *G* = (*S ∪ A, E*), a labelled bipartite graph with *N* vertices in one part *S*, and *M* vertices in the other part *A*. The vertices in *S* denote the strains and the vertices in *A* denote the antibiotics. All vertices in *S* are connected to all vertices in *A*, but the (labelled) edges may have different weights. In particular, the label of the edge going from a vertex *i ∈ S* to a vertex *j ∈ A* is exactly the element *T*_*ij*_. In other words, the edge labels denote the phenotype of strain *i* with respect to antibiotic *j*. See Figure A.3 for a depiction of this operation.

**Fig. A.3:**
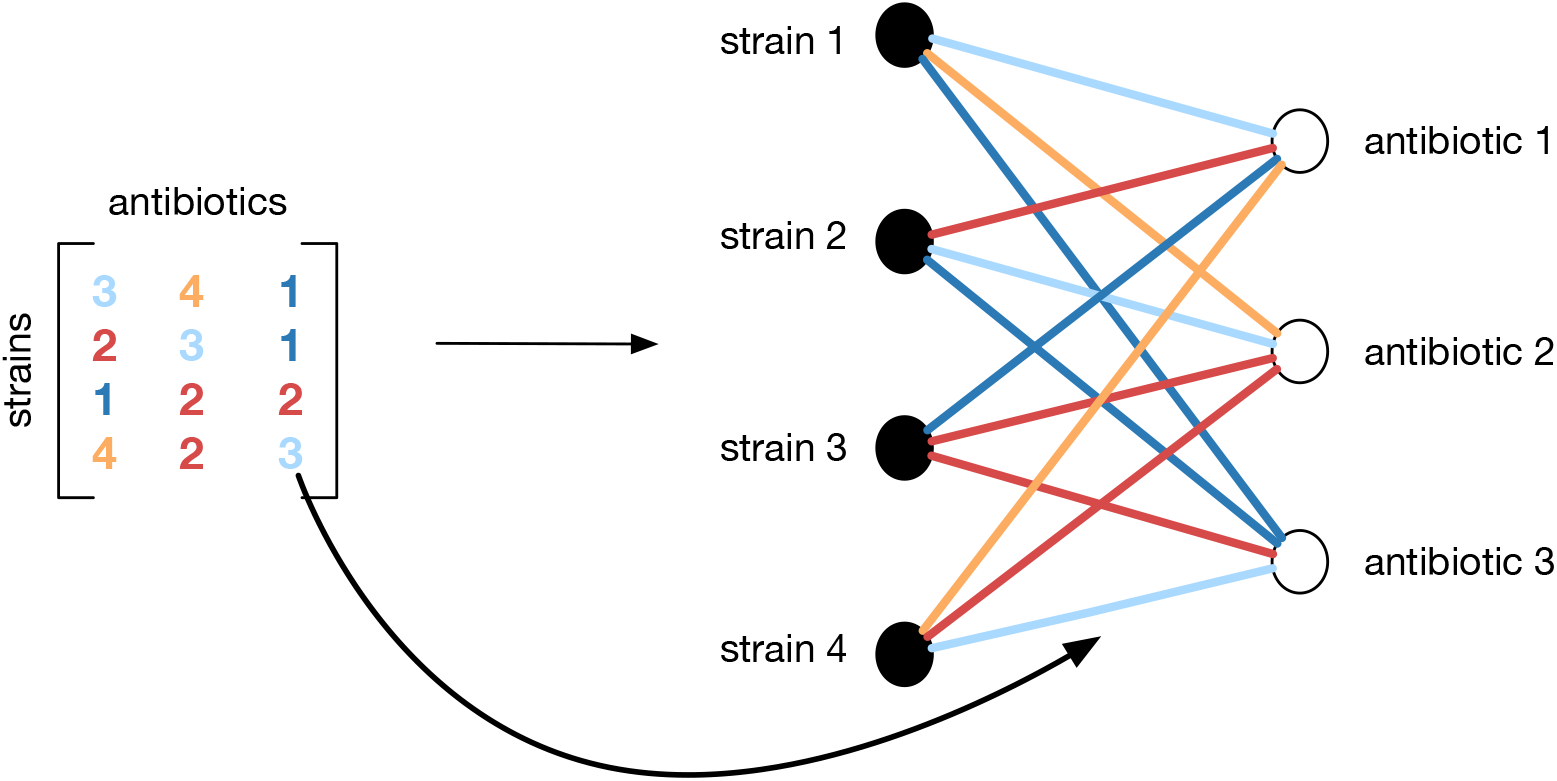
This invariance of the community under relabelling is exactly what is captured by a graph - the community is described only by the phenotype of each strain with respect to each antibiotic. We envision the matrices in 𝕋_*N,M*_ as adjacency matrices of labelled, bipartite graphs, and claim that the graphs i.e., the relative connections of strains and antibiotics, uniquely specifies a community, *but only upto isomorphism*. This figure shows an example of the transformation from 𝕋_4,3_ to 𝔾_4,3_, which is defined here and also for any *N, M* as the image of 𝕋_4,3_ under this map given by interpreting the matrices as specifying the edges of a graph. Since there are efficient algorithms to check for isomorphisms between graphs, we develop an algorithm to derive a set of non-isomorphic graphs instead of manipulating the matrices themselves, which is in principle possible. In other words, we want a set of graphs such that given any matrix in𝕋_*N,M*_ for some *N* and *M*, there is a graph in our collection that corresponds to this matrix, perhaps after relabelling strains and antibiotics.

An *isomorphism* between 2 graphs *X* and *Y* is a one-to-one correspondence *f* between the vertex sets of *X* and *Y* such that any two vertices *u* and *v* of *X* are adjacent in *X* if and only if *f* (*u*) and *f* (*v*) are adjacent in *Y*, with the edges (*u, v*) and (*f* (*u*), *f* (*v*)) having same edge weights and direction. Two graphs are called isomorphic to each other if there exists an isomorphism between them. See Figure

A.4 for an example of two isomorphic graphs. Intuitively, an isomorphism corresponds to a relabelling of the vertices or a re-drawing of the graph. The *isomorphism class* of a graph is the set of all graphs it is isomorphic to. Importantly, note that any two isomorphic graphs have the same isomorphism class.

**Fig. A.4:**
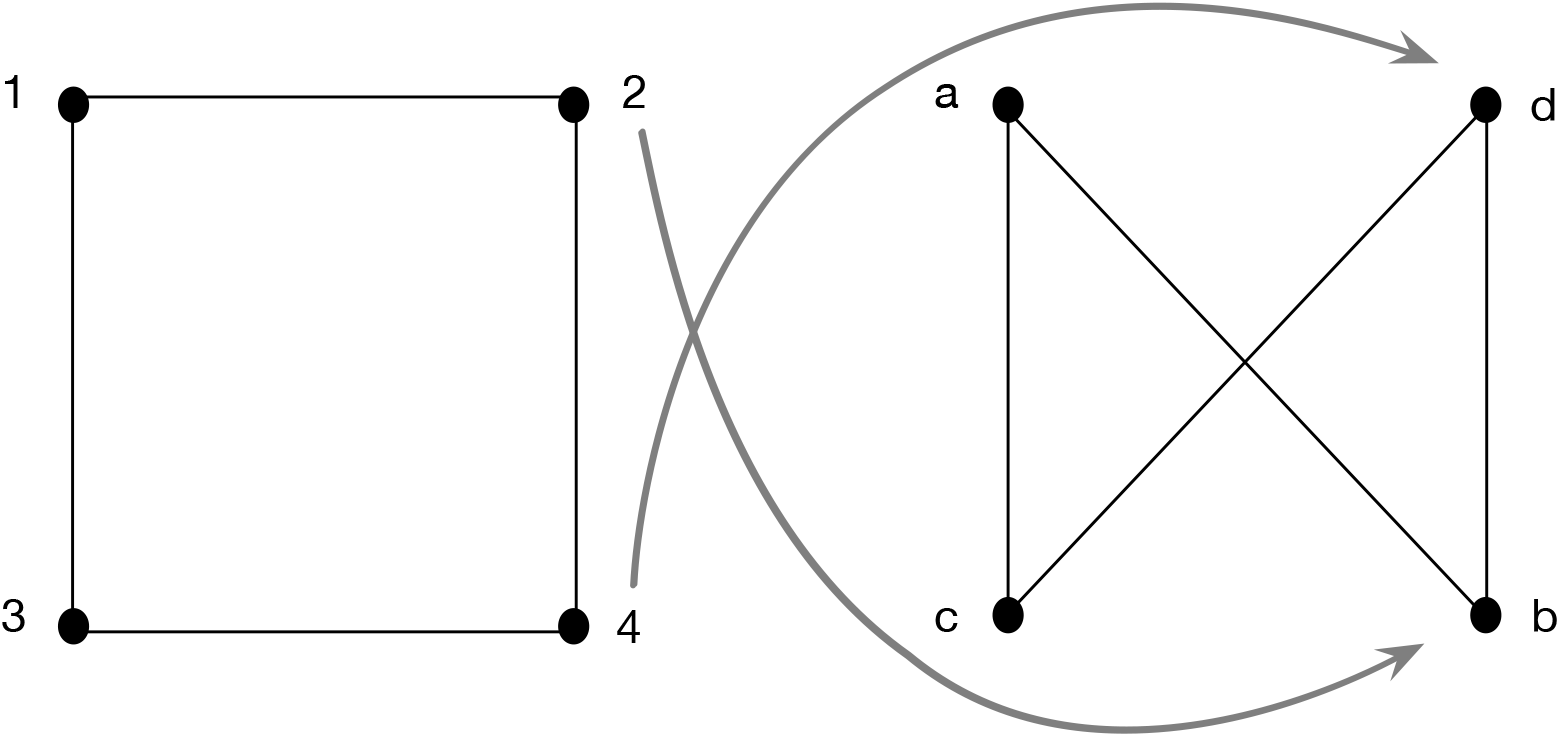
Two graphs are isomorphic if they are just re-drawings of each other - that is, if there is a correspondence of their vertices such that they look the same. This figure shows an example of two isomorphic graphs on 4 vertices. The isomorphism is as follows: 1 → *a*, 2 → *b*, 3 → *c*, 4 → *d*. Intuitively, the graph on the right is a re-drawing of that on the left since it can be obtained by twisting the right side of the square (and thus the horizontal edges as well) when one physically “flips” vertices 2 and 4 out of the plane of the page.

The remaining paragraphs describe the method by which the reduction is performed on G_*M,N*_. We claim that it is sufficient to only consider one element from the isomorphism class of each graph in G_*M,N*_. The redundancy is hence removed by discarding graphs which fall into the isomorphism class of a graph already considered, since isomorphic graphs have the same isomorphism class. The procedure can be described as follows:

- Initialize a storage structure *S*.
- Generate all matrices in T_*N,M*_.
- Now we iterate over 𝕋_*N,M*_ - for each matrix *T*, convert *T* to the corresponding bipartite graph *G* as above.
- For each graph already present in *S*, check if it is isomorphic to *G*.
- *G* is discarded as soon as we encounter a graph in *S* that is isomorphic to it.
- In other words, *G* is added to *S* iff it is non-isomorphic to all graphs already present in *S*.

All that remains is to prove that this procedure gives rise to a subset of 𝔾_*M,N*_ which is in one-to-one correspondence with the set Ω_*N,M*_ of all communities with *N* strains and *M* antibiotics. Recall that 𝕋_*N,M*_ already describes Ω_*N,M*_ fully, but redundantly. By our construction of *S*, we see that no pair of graphs in *S* can be isomorphic to each other. Therefore, we need to prove that two antibiotic profiles describe the same community in Ω_*N,M*_ if and only if they are isomorphic.

### Remark

For the more mathematically inclined, this “reduction” amounts to interpreting the matrices in 𝕋_*M,N*_ as adjacency matrices of the graphs in 𝔾_*M,N*_ and taking the quotient of 𝔾_*M,N*_ by the equivalence relation induced by graph isomorphism. Then we map the non-redundant set of graphs back to their adjacency matrices – this map would have failed to be injective if the redundant graphs were not removed.

First suppose we construct two antibiotic profiles *X, Y* for the same community by adopting two (possibly different) labellings. In this case, we can explicitly construct an isomorphism between *X* and *Y* since we know the labels of each strain in both *X* and *Y*. Conversely, suppose we have two isomorphic antibiotic profiles. Both must necessarily be bipartite. We can match the strain parts and the antibiotic parts of the two graphs by looking at the vertex labels - the strain parts are the parts which have vertices labelled ‘S’. We can then consult the isomorphism to infer which vertices are mapped to each other. By definition of an isomorphism, the mapped vertices will have the same edges and hence the same phenotype. This completes the proof.

### Remark

The above if and only if condition also means that each matrix in the output of this algorithm represents a distinct community. In other words, our algorithm performs the maximum possible reduction in search space.

## A.2 Detailed stochastic spatial model

Suppose there are *N* strains and *M* antibiotics. We initialise an *L* × *L* lattice and set the initial condition for population dynamics as follows: for each grid cell, a “spore” of type *i* (*i* = 1, …, *N*) is placed at this grid cell with probability *p*, where *p* is chosen such that *Np* < 1. These spores then reproduce and spread over the lattice to occupy all grid cells. We drop the assumption that individuals can disperse over large distances, which differentiates this model from the well-mixed model of previous sections. One *time step* or *generation* is defined as the total time taken to update the state of each individual currently on the lattice. An *update step* is defined as the process of updating the state of a single individual.

At each time step, we iterate over all grid cells on the lattice sequentially. If the grid cell is empty, we move on to the next. If the grid cell is occupied by an individual A, there are two processes taking place - first, consider the diffusion of antibiotics and degrader chemicals. We depart from the simplifying assumption made by Kelsic et al. (2015) that antibiotics and degraders are binary in their effect, i.e., either fully effective within an area of *K*_*P*_ or *K*_*D*_ or not effective at all outside this area [1]. We assume each individual produces a fixed amount of antibiotic/degrader chemicals per update step. If A produces any antibiotics, then *u*_*p*_ units of each antibiotic type are placed on the focal grid cell. Similarly, if A degrades any antibiotics, *u*_*d*_ units of each degrader chemical are placed on all neighbouring grid cells and diffused as above. Once all chemicals have been secreted, a Gaussian filter is applied independently to the concentrations of each antibiotic to simulate their diffusion. On a grid cell with a non-zero amount of antibiotic and corresponding degrader, we assume that the resultant antibiotic amount, i.e., post-inactivation by degradation, is the difference between the number of antibiotic and degrader units present in this cell. This is equivalent to imposing that one unit of the degrader chemical “reacts” with precisely one unit of the antibiotic. There might indeed be different stoichiometries in natural communities. Still, this condition allows us to make conclusions directly about the interaction graphs without any confounding effects due to the reactions’ dynamics.

Next, we consider the birth and death of individuals: The growth rates of individuals of a given phenotype are decided using the method followed in the previous section. Following Vetsigian [2], the effect of the antibiotics on the sensitive strains is decided by a dose-response curve with a threshold dosage below which antibiotics have no effect. Suppose the antibiotic concentration of any antibiotic (that A is sensitive to) is above the threshold. In that case, A is killed with some probability, a function of the antibiotic concentration. The dose-response curve gives the probability of dying as a function of external antibiotic concentration:

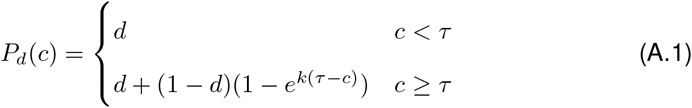

where the constants *τ* and *k* are to be determined. Respectively, they denote the threshold concentration value and the susceptibility of the sensitive strain i.e., how quickly the probability of death asymptotes to 1. These constants are determined by imposing two constraints on *P*_*d*_(*c*). Let *G*(*r*) denote the value of the 2-dimensional Gaussian at a distance *r* from the peak (recall that it is spherically symmetric). Since *u*_*p*_ is the volume of antibiotic secreted by a producer every time step, *u*_*p*_*G*(*r*) is the concentration, post one application of the Gaussian filter, at a distance *r*. Then the conditions imposed are

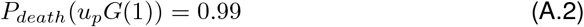

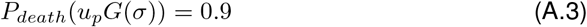

These constraints correspond to a “common sense” expectation from the dose-response curve. The first constraint is motivated by the expectation that the death probability must be very high (but perhaps not 1) for all neighbours of the producer i.e., all individuals at distance 1. The second corresponds to the expectation that at a distance of *σ* (here *σ* > 1), the death probability must be high, but lesser than the death probability at distance 1. Let

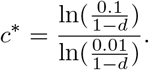

Solving the two constraints for *τ* and *k* simultaneously, we have

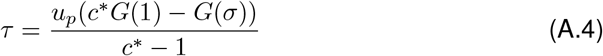

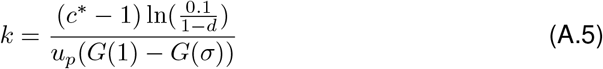

The width *σ* of the Gaussian filter is set as *K*_*P*_/3. This is because *K*_*P*_ in the mixed-inhibition zone model was interpreted as the area outside which the antibiotic does not have an effect. We assume the area of the Gaussian inside a distance of 3*σ* is an approximation of this area of effect since it contains 99.7% of the area under the normal distribution curve.

For the dispersal of offspring, we consider a dispersal neighbourhood D consisting of some arrangement of grid cells neighbouring the cell that A occupies. If any of these cells are empty, then another individual belonging to the same strain as A is born with probability *r*, a function of the growth rate of A and the antibiotic concentration on that grid cell. Then this new individual is placed at a grid cell picked uniformly from the empty cells in D.

This simulation is then run for many generations, one would then check for communities in which the strain abundances are nonzero for a long time. In particular, we study the 1-antibiotic PSD motif and simulate its dynamics over a large parameter range. To compare the dynamics under different parameter values after the system has relaxed to an attractor, if it exists, we take the average of the abundance over the last 500 generations, where the simulation was run for a total of 6000 generations. The implicit assumption we are making here is that the attractor will be reached after 6000 generations, and this generally seems to be true in our simulations.

## Footnotes

1 It is not helpful to consider *N* > 4 since for any *M*, there can be at most 4^*M*^ distinct phenotypes. Thus when *N* > 4^*M*^, there will be at least two strains with the same phenotype, thereby reducing this case to a system of equations that we have already analysed.

2 Note that some communities have uncountably many stable fixed points which disappear entirely if the two antibiotics have different metabolic costs. The assumption of identical metabolic costs is made only for convenience and does not represent most communities, so we omit such cases.

